# The phosphatase Bph and peptidyl-prolyl isomerase PrsA are required for gelatinase expression and activity in *Enterococcus faecalis*

**DOI:** 10.1101/2022.04.06.487426

**Authors:** Julia L. E. Willett, Ethan B. Robertson, Gary M. Dunny

## Abstract

*Enterococcus faecalis* is a common commensal bacterium in the gastrointestinal tract as well as a frequent nosocomial pathogen. The secreted metalloprotease gelatinase (GelE) is an important *E. faecalis* virulence factor that contributes to numerous cellular activities such as autolysis, biofilm formation, and biofilm-associated antibiotic resistance. Expression of *gelE* has been extensively studied and is regulated by the Fsr quorum-sensing system. Here, we identify two additional factors regulating gelatinase expression and activity in *E. faecalis* OG1RF. The Bph phosphatase is required for expression of *gelE* in an Fsr-dependent manner. Additionally, the membrane-anchored protein foldase PrsA is required for GelE activity, but not *fsr* or *gelE* gene expression. Disrupting *prsA* also leads to increased antibiotic sensitivity in biofilms independent of the loss of GelE activity. Together, our results expand the model for gelatinase production in *E. faecalis*, which has important implications for fundamental studies of GelE function in *Enterococcus* and also *E. faecalis* pathogenesis.

**Importance:** In *Enterococcus faecalis*, gelatinase (GelE) is a virulence factor that is also important for biofilm formation and interactions with other microbes as well as the host immune system. The long-standing model for GelE production is that the Fsr quorum sensing system positively regulates expression of *gelE*. Here, we update that model by identifying two additional factors that contribute to gelatinase production. The biofilm-associated Bph phosphatase regulates the expression of *gelE* through Fsr, and the peptidyl-prolyl isomerase PrsA is required for production of active GelE through an Fsr-independent mechanism. This provides important insight into how regulatory networks outside of the *fsr* locus coordinate expression of gelatinase.

## Introduction

*Enterococcus faecalis* is a commensal bacterium in the gastrointestinal tracts of hosts ranging from insects to humans (1). It is also a prevalent human pathogen that causes biofilm-associated infections such as endocarditis, urinary tract infections, and infections at wounds and surgical sites (2, 3). A major virulence factor for *E. faecalis* is gelatinase (GelE), a secreted zinc metalloprotease that mediates chain length and autolysis as well as host intestinal epithelial permeability (4–6). GelE also processes numerous substrates, including the *Candida albicans*-inhibiting peptide EntV (EF1097, enterocin O16) (7, 8), the major *E. faecalis* autolysin AtlA (9), pheromone peptides that induce conjugative plasmid transfer in *E. faecalis* (5, 10), and a component of the enterohemorrhagic *E. coli* type 3 secretion system (11). Expression of *gelE* is positively regulated by the well-studied Fsr quorum sensing system, which is encoded by the *fsrABDC* locus (12–14). Genotypic and phenotypic screens of *E. faecalis* clinical isolates have identified mutations that abrogate production of GelE, including a 23.9 kb deletion encompassing the *fsr* locus as well as truncations and IS256 insertions in *fsrC* (15–20).

Here, we identified two additional gene products that modulate gelatinase expression and activity in *E. faecalis* OG1RF. Using RNAseq, we found that deletion of the gene encoding the biofilm-associated phosphatase Bph (Δ*bph*) decreases expression of the *fsr* locus and thus *gelE*. Δ*bph* also had decreased expression of an uncharacterized locus encoding multiple cell surface WxL-domain proteins, which contributes to *in vitro* biofilm formation. Through separate experiments, we found that the peptidyl-prolyl isomerase (PPIase) PrsA is required for GelE activity in an Fsr-independent manner, and we hypothesize that PrsA is required for correct folding (and therefore activity) of secreted GelE. PrsA was previously linked to salt tolerance and virulence in *E. faecalis* (21). Additionally, PrsA homologs in other Gram-positive bacteria are required for activity of secreted proteins (22–24) and resistance to oxacillin and vancomycin (25, 26). We observed a similar pattern in OG1RF, although increased antibiotic sensitivity was detected in biofilms and colony growth on plates, but not liquid culture. These results describe two new regulators of the virulence factor gelatinase, highlight the global effects of disrupting *bph* and *prsA* in *E. faecalis*, and provide insight into biofilm-specific responses to antibiotics when secreted proteins are misfolded.

## Results

### Deletion of *bph* decreases expression of *gelE* through the Fsr quorum sensing system

We previously reported that *E. faecalis* Bph is a phosphatase required for biofilm formation and found widespread differences in protein expression in a Δ*bph* mutant compared to the parental OG1RF strain (27). To determine whether these changes were due to differential gene expression, we used RNAseq to compare transcripts present in OG1RF and Δ*bph* after 2 and 4 h planktonic growth. Using a significance cutoff of q<0.05 and a log_2_-fold change cutoff of +/-2, we identified 133 differentially expressed transcripts in Δ*bph* at 2 h (42 upregulated, 91 downregulated) and 218 differentially expressed transcripts at 4 h (119 upregulated, 99 downregulated) (**Supplementary Table 1**, Figure S1). Differentially expressed genes included many involved in carbohydrate/nucleotide metabolism, membrane transport, peptidoglycan biosynthesis, ribosomal protein synthesis, protein secretion, and DNA replication and repair. Additionally, the ethanolamine utilization operon (OG1RF_11333-11351) plus genes involved in selenium and molybdenum metabolism and folate biosynthesis (OG1RF_11941-11962 (28) and OG1RF_11179-11189) were significantly upregulated at 4 h (**Supplementary Table 1**). Curiously, *xdh* (OG1RF_11951, xanthine dehydrogenase) is upregulated in biofilms (29), although the increased expression of *xdh* in Δ*bph* does not correlate with increased biofilm production.

Surprisingly, expression of the Fsr quorum sensing system as well as the Fsr-regulated proteases *gelE* and *sprE* was significantly reduced in Δ*bph* compared to OG1RF (Figure 1A, **Supplementary Table 1**), although we previously found that Δ*bph* has a gelatinase (GelE)-positive phenotype on gelatin plates (27). Because *fsr*, *gelE*, and *sprE* expression increases with greater cell density, we used qRT-PCR to measure *gelE* expression in Δ*bph* at 6 and 8 h. We measured a ∼92% decrease in transcript levels relative to OG1RF at 6 hr and a ∼95% decrease at 8 hr, (Figure 1B), suggesting that the Fsr system is downregulated throughout Δ*bph* growth.

**Figure 1:**
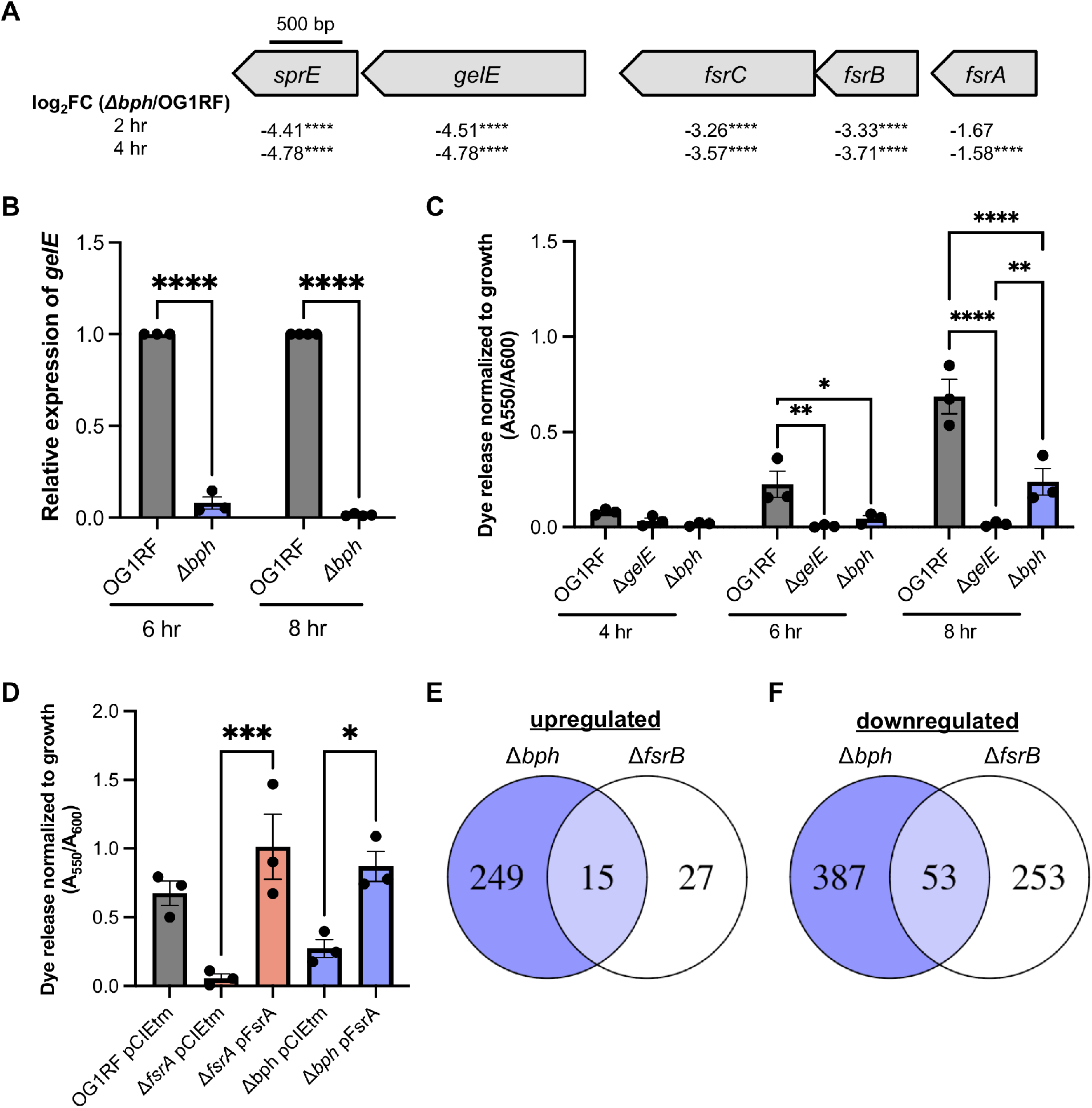
Deletion of *bph* reduces *gelE* expression and GelE activity through reduced expression of the Fsr quorum sensing system. **A)** Genetic organization of *fsrABC*, *gelE*, and *sprE* and log_2_FC values from RNAseq at 2 and 4 h. ****p<0.0001. **B)** Relative levels of *gelE* transcripts in OG1RF and Δ*bph* at 6 and 8 h as measured by qRT-PCR. Three biological replicates were tested at 6 h, and 4 biological replicates were tested at 8 h. ****p<0.0001 by two-way ANOVA with Sidak’s test for multiple comparisons. **C)** Quantification of dye release (A_550_) from OG1RF, Δ*gelE*, and Δ*bph* supernatants relative to growth (A_600_) at 4, 6, and 8 h. Three biological replicates were tested for each strain. **p<0.01, ***p<0.001 by two-way ANOVA with Sidak’s test for multiple comparisons. **D)** Rescue of gelatinase activity in Δ*bph* by expression of *fsrA* from a pheromone-inducible plasmid. Three biological replicates were tested for all strains. *p<0.05, ***p<0.001 by one-way ANOVA with Sidak’s test for multiple comparisons. For panels **B**, **C**, and **D**, each data point represents an independent biological replicate, and error bars represent standard error of the mean. **E, F**) Venn diagrams comparing upregulated and downregulated genes in Δ*bph* and Δ*fsrB* mutants.

We next quantified GelE activity using dye release assays to resolve the incongruent expression and phenotype in the Δ*bph* mutant. Gelatin plate-based assays show cumulative GelE activity after overnight colony growth and may obscure subtle phenotypes. We isolated supernatants from OG1RF, Δ*gelE*, and Δ*bph* cultures at 4, 6, and 8 h and mixed them with the protease substrate Azocoll. Proteases like GelE degrade the insoluble dye-protein complex and release soluble Azo dyes, which can be quantified by measuring absorbance (30). Dye released by OG1RF supernatants increased over time, while no dye release was detected in the Δ*gelE* supernatants (Figure 1C). Dye release from Δ*bph* supernatants decreased 70-80% relative to OG1RF at each time point. Gelatinase activity in Δ*bph* was significantly increased relative to Δ*gelE* at 8 h but not 4 or 6 h. This suggests that despite reduced *gelE* expression in Δ*bph*, low levels of active GelE accumulate over time, resulting in the GelE-positive phenotype previously observed after overnight growth on plates (27).

Low levels of *fsrA* transcript in Δ*bph* could be caused by reduced gene expression or RNA degradation. We reasoned that if transcription of *fsrA* was reduced, then expressing *fsrA* from a plasmid could restore GelE activity. However, if *fsrA* transcripts were degraded in the Δ*bph* background, then plasmid-borne *fsrA* would not restore GelE activity. We cloned *fsrA* into a pheromone-inducible backbone, induced expression in Δ*fsrA* and Δ*bph*, and quantified GelE activity in the supernatants using Azocoll assays. In both strains, expression of *fsrA* from a plasmid significantly increased GelE activity relative to the empty vector control (Figure 1D, 17.8-fold and 3.19-fold, respectively). Overall, we conclude that Bph regulates *gelE* expression through Fsr and that additional regulatory networks outside of Fsr are disrupted in the absence of *bph*.

### Comparison of differentially expressed genes in Δ*bph* and Δ*fsrB* deletion mutants

Next, we asked how much of the altered gene expression in the Δ*bph* mutant could be linked to disruption of the Fsr quorum sensing system. We compared all differentially expressed genes identified using RNAseq to published microarray data from a Δ*fsrB* mutant (31). In Δ*fsrB*, 15 of the 42 upregulated genes were shared with Δ*bph*, and 53 of the 306 downregulated genes were shared with Δ*bph* (Figure 1EF). Of the 15 common upregulated genes, 8 are within the ethanolamine utilization operon (**Supplementary Table 2**). Shared downregulated genes included 5 genes predicted to encode an uncharacterized surface protein complex (OG1RF_10485-10489, OG1RF_10491). To determine whether reduced expression of OG1RF_10485-10491 contributes to the biofilm-deficient phenotype of Δ*bph* (27), we obtained Tn mutants from an arrayed Tn library (32) and measured biofilm production in 96-well plates. OG1RF_10490::Tn had reduced biofilm relative to OG1RF at 6 h and 24 h, and OG1RF_10489::Tn and OG1RF_10492::Tn had reduced biofilm at 24 h (Figure S2). Interestingly, 4 genes in the enterococcal polysaccharide antigen (Epa) operon (OG1RF_11721-11724) were also downregulated in both Δ*bph* and Δ*fsrA*, although we did not previously observe changes associated with altered Epa synthesis in polysaccharides purified from Δ*bph* (27, 33, 34).

### The peptidyl-prolyl isomerase PrsA is required for GelE functionality independent of Fsr

In a previous screen for Tn mutants with defects in biofilm formation, we found that a mutant with a Tn insertion in *prsA* (OG1RF_10423) had a GelE-negative phenotype on gelatin plates (35). PrsA is an extracellular parvulin-like peptidyl-prolyl isomerase (PPIase) (Figure 2A) that is required for activity of secreted proteins and membrane proteins in numerous Gram-positive bacteria, including *Bacillus anthracis* (23), *Bacillus subtilis* (36), *Listeria monocytogenes* (24), *Staphylococcus aureus* (37), group A *Streptococcus* (38), and *Streptococcus equi* (39). In *E. faecalis*, PrsA is important for survival in high salt concentrations and virulence in *G. mellonella* (21), although no role in folding specific protein substrates has been described. We confirmed the gelatinase-negative phenotype of the *prsA* Tn mutant (*prsA*::Tn) grown in both TSB-D and MM9-YEG growth media, both of which were previously used when studying biofilm formation by *prsA*::Tn (Figure 2B). Expression of *prsA* from a pheromone-inducible plasmid restored GelE activity on plates (Figure 2B). We next measured GelE activity using dye-release assays with supernatants from cells cultured for 6 h. Relative to OG1RF, dye release from *prsA*::Tn supernatants was reduced approximately 70% in both media, and dye release was restored in *prsA*::Tn by expression of *prsA* from a plasmid (Figure 2C).

**Figure 2.**
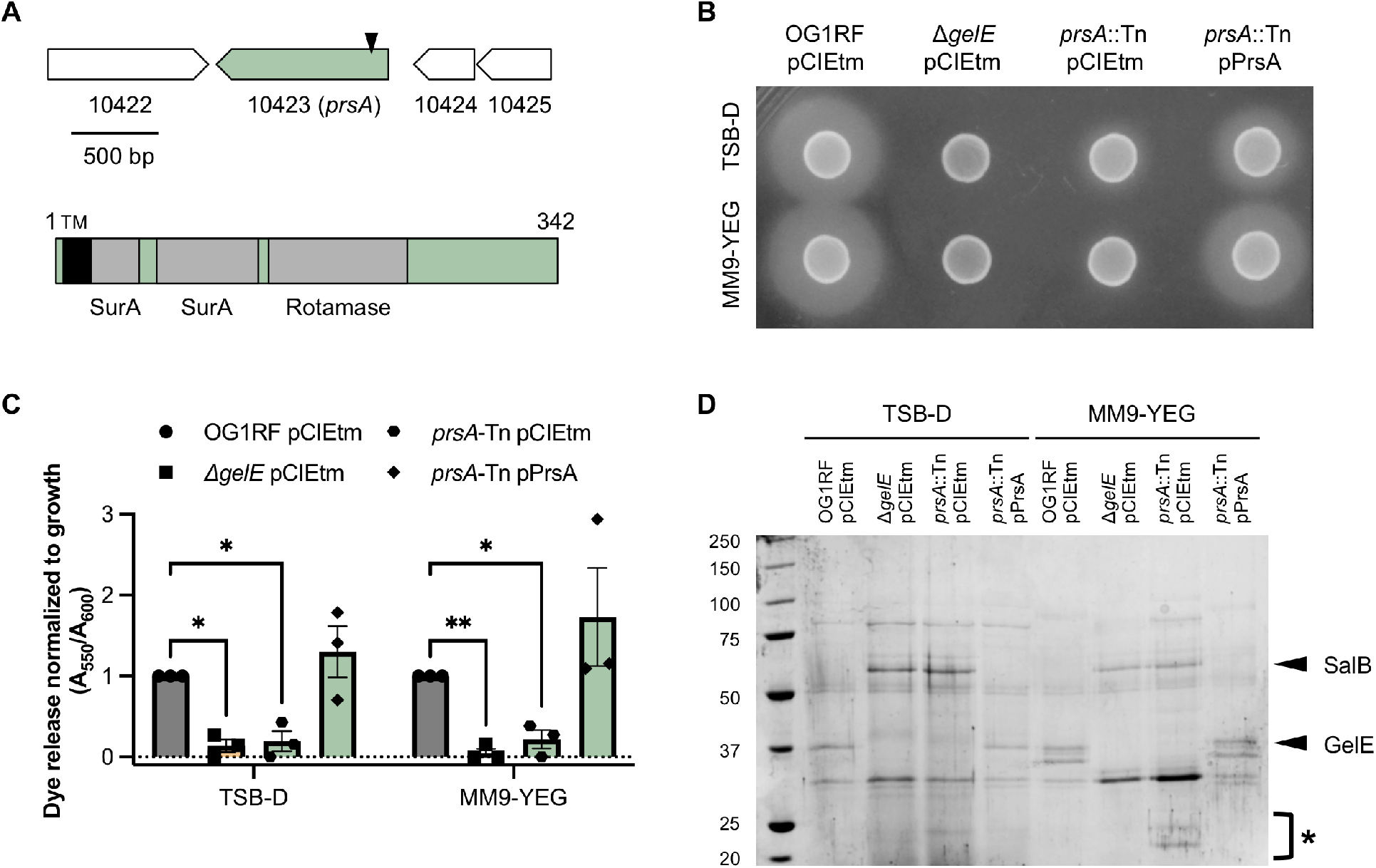
PrsA is a predicted extracellular peptidyl-prolyl isomerase that is required for gelatinase functionality in OG1RF. **A)** Cartoon (top) showing *prsA* and the location of the Tn insertion (black triangle) and PrsA protein features (bottom). Pfam domains, gray rectangle; transmembrane domain, black rectangle. **B)** Gelatinase plate assay with OG1RF pCIEtm, Δ*gelE* pCIEtm and *prsA*::Tn pCIEtm, and *prsA*::Tn pPrsA. Hazy zones around colonies are indicative of gelatinase activity. Image is representative of three biological replicates. **C)** Quantification of relative gelatinase activity compared to OG1RF using an Azocoll dye-release assay. Three biological replicates were tested for each strain. *p<0.05, **p<0.01 by two-way ANOVA with Dunnett’s test for multiple comparisons. **D)** SDS-PAGE evaluation of exoproteins using the same strains as **B** and **C**. Prominent secreted proteins (SalB, GelE) are marked with arrows, and small protein bands in *prsA*::Tn are marked with *. Image is representative of three biological replicates.

Given that GelE processes secreted proteins (5, 40), a loss of GelE activity in the *prsA* mutant background could also affect other supernatant proteins. We used SDS-PAGE to evaluate supernatant proteins from cells grown in TSB-D and MM9-YEG and found that the *prsA*::Tn supernatant lacked the prominent GelE band (41) (Figure 2D). The Δ*gelE* and *prsA*::Tn supernatants had similar protein banding patterns, and additional small protein bands were visible in the *prsA*::Tn samples (Figure 2D, marked with *). Previous work showed the Δ*gelE* mutant had an increase in supernatant levels of the secreted protein SalB, which contributes to cell envelope integrity and cephalosporin resistance (40, 42, 43). Additionally, GelE processes recombinant SalB (43). We observed a band corresponding to SalB in the *prsA*::Tn supernatants (Figure 2D). SalB levels are also increased in Δ*bph* supernatants (27), so decreased GelE levels in the Δ*bph* and *prsA*::Tn supernatants likely lead to increased accumulation of SalB. Expression of *prsA* from a plasmid restored *prsA*::Tn supernatant proteins to those of parental OG1RF. We also observed minor differences in protein banding patterns between OG1RF and *prsA*::Tn in protoplast and cell wall samples (Figure S3). Together, these experiments demonstrate that disruption of *prsA* abrogates gelatinase production and alters the exoprotein profile of OG1RF.

Because *gelE* expression is reduced in Δ*bph*, we also considered that GelE levels in *prsA*::Tn could be reduced due to changes in gene expression. We examined expression from *fsrA*, *fsrB*, and *gelE* promoters using *lacZ* promoter fusion constructs and found no significant differences in expression between *prsA*::Tn and OG1RF (Figure 3AB), suggesting that reduced GelE levels in *prsA*::Tn are not due to reduced transcription. We then asked whether the gelatinase-negative phenotypes of Δ*bph* and *prsA*::Tn could be rescued by expression of *gelE* from a plasmid. We hypothesized that if the gelatinase-negative phenotype of *prsA*::Tn resulted from aberrant folding of GelE, then expression *in trans* would not rescue the phenotype in Azocoll assays. Nisin induction of pMSP3535::*gelE* significantly increased dye release from Δ*gelE* and Δ*bph* supernatants (21.0-fold and 3.0-fold, respectively), but not from *prsA*::Tn (Figure 3C). This demonstrates that the loss of GelE activity in *prsA*::Tn is not caused by changes in *fsr* or *gelE* gene expression.

**Figure 3.**
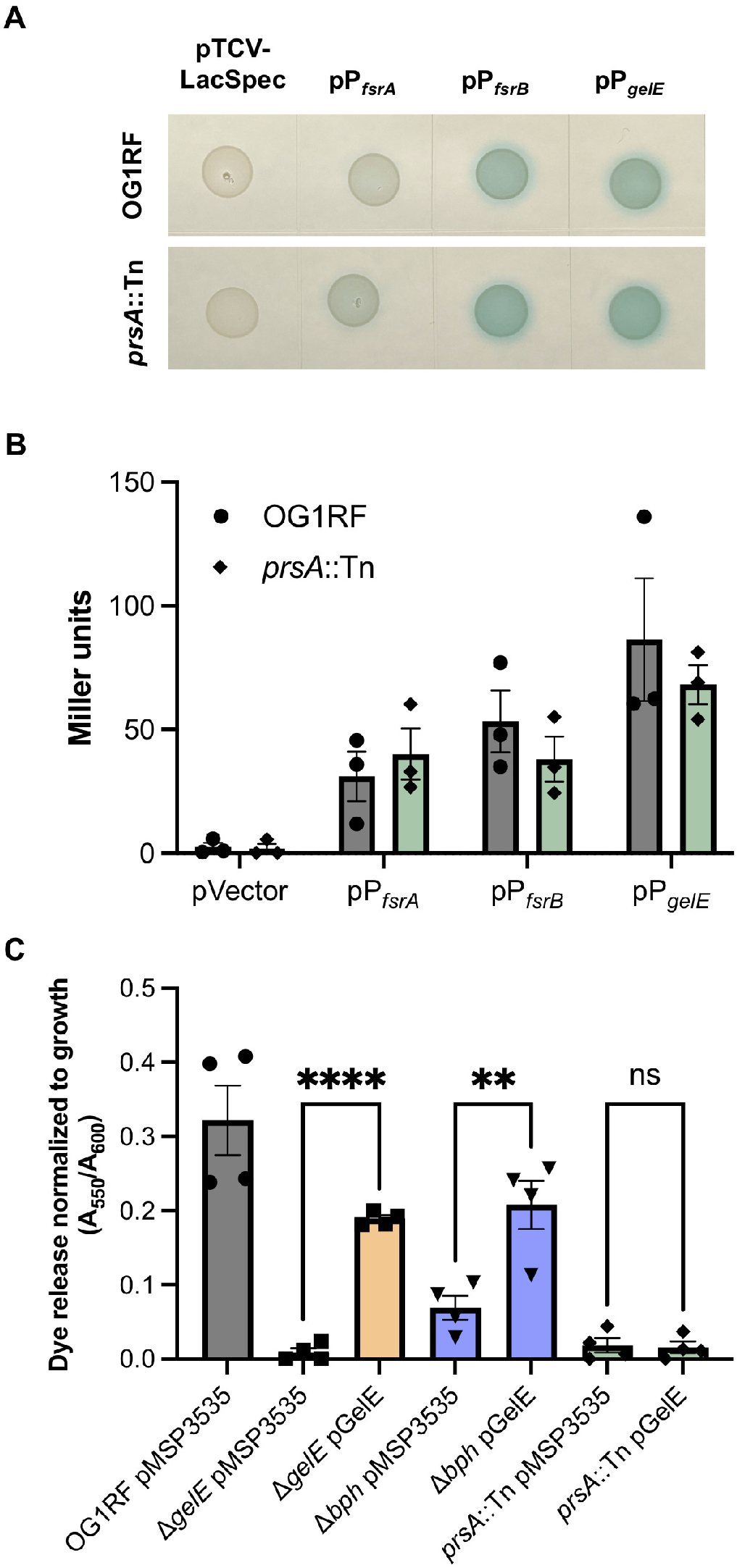
Loss of GelE in *prsA*::Tn is not due to reduced expression of the Fsr quorum sensing system. **A)** Empty pTCV-LacSpec or derivatives carrying the promoter regions of *fsrA*, *fsrB*, or *gelE* were transformed into OG1RF and *prsA*::Tn and spotted on MM9-YEG agar plates supplemented with 5-bromo-4-chloro-3-indolyl-β-D-galactopyranoside (X-gal). Images are representative of 3 independent experiments. **B)** Promoter activities of the strains from panel A were quantified with β-galactosidase assays (n = 3). **C)** Expression of *gelE* from a pheromone-inducible plasmid (pMSP3614) rescues gelatinase activity in Δ*gelE* and Δ*bph* but not *prsA*::Tn (n = 4). In panels **B** and **C**, each data point represents a biological replicate. **p<0.01, ****p<0.0001 by one-way ANOVA with Sidak’s multiple comparisons test.

### PrsA mediates biofilm-associated antibiotic resistance independent of GelE

Deletion of *prsA* in *S. aureus* isolates leads to increased sensitivity to glycopeptides and oxacillin (25, 26), and disruption of *prsA* in *B. subtilis* results in destabilized penicillin-binding proteins (36). However, in *E. faecalis* V583, deletion of *prsA* did not alter susceptibility to antibiotics during growth on agar plates (21). Therefore, we wondered whether disrupting *prsA* in the OG1RF background would lead to altered susceptibility to cell wall-active antibiotics. We measured growth of OG1RF, Δ*gelE*, Δ*bph*, and *prsA*::Tn in 2-fold dilution series of ampicillin, oxacillin, penicillin G, and vancomycin relative to untreated cultures.

In liquid culture, no change in sensitivity was observed for *prsA*::Tn in oxacillin, penicillin G, or vancomycin (Figure 4A-C). *prsA*::Tn was slightly more susceptible than the other strains to 8 µg/mL ampicillin (Figure 4D). Interestingly, Δ*gelE* grew to a higher OD_600_ in sub-inhibitory antibiotic concentrations. Next, we spotted serial dilutions of each strain onto agar plates supplemented with sub-inhibitory concentrations of each antibiotic (empirically chosen based on the liquid growth assays). Surprisingly, growth of *prsA*::Tn but not the Δ*gelE* strain was strongly inhibited relative to OG1RF on plates containing 2 µg/mL oxacillin or vancomycin (Figure 4E), even though no growth defect was observed in liquid culture for either antibiotic at this concentration. None of the mutant strains had growth defects on plates containing sub-inhibitory concentrations of ampicillin or penicillin G (Figure 4E).

**Figure 4.**
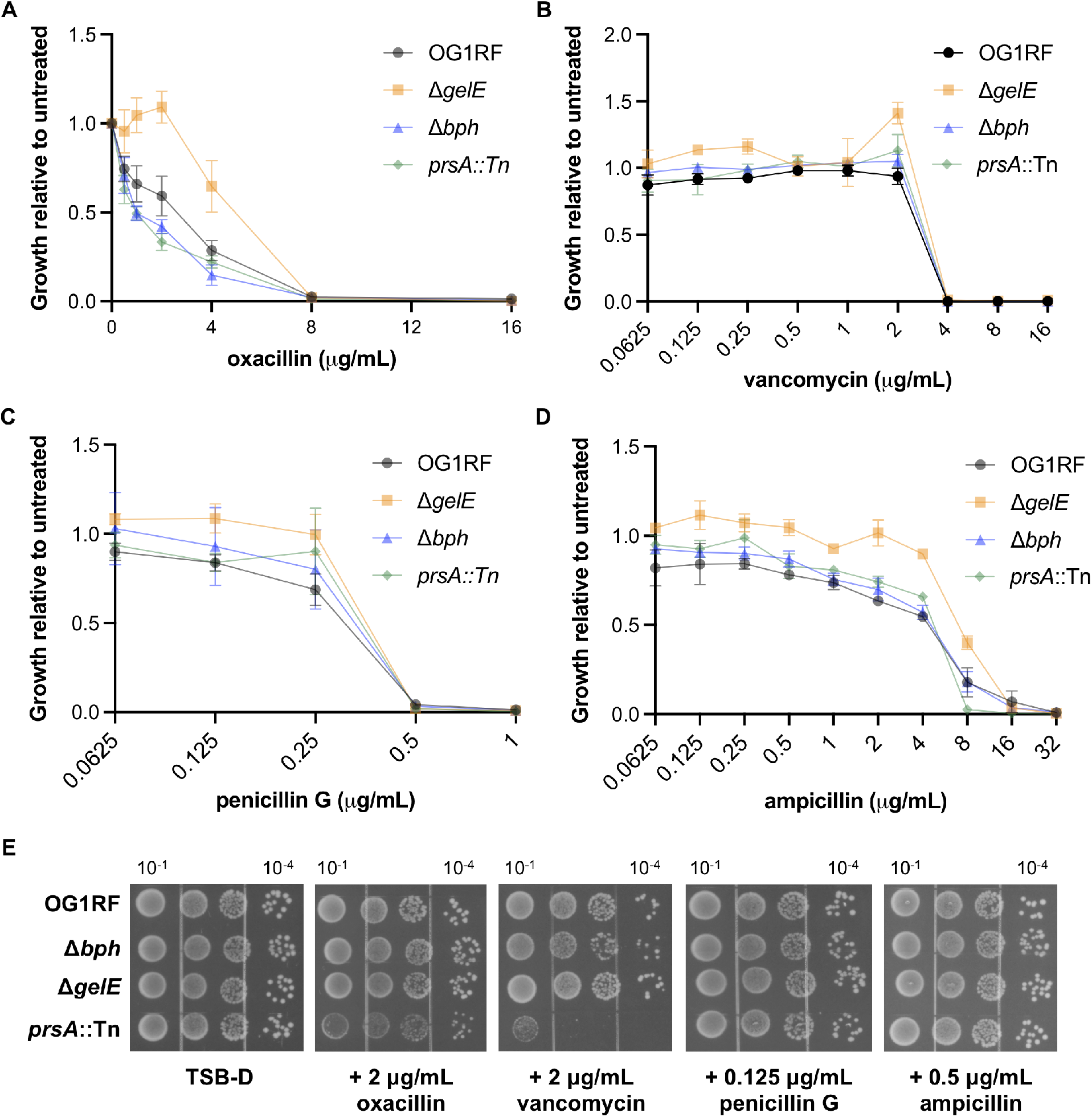
Growth of OG1RF, Δ*gelE*, Δ*bph*, and *prsA*::Tn in antibiotics that target the cell envelope. The indicated strains were grown in TSB-D in a 2-fold dilution series of **A)** oxacillin, **B)** vancomycin, **C)** penicillin G, and **D)** ampicillin. Growth at each concentration was calculated relative to untreated samples. Data points represent the average of 3 biological replicates, and error bars show standard error of the mean. **E)** Overnight cultures were normalized to 10^7^ CFU/mL and serially diluted. Dilutions (shown above plate images) were spotted onto TSB-D plates supplemented with the indicated sub-inhibitory concentrations of antibiotics. Gel images are representative of 3 biological replicates.

We previously reported that OG1RF mutants with Tn insertions in *fsrA* and *gelE* had decreased biofilm formation in the presence of sub-inhibitory concentrations of daptomycin, gentamicin, and linezolid (34). Therefore, we wondered whether reduced levels of GelE in our mutant strains would lead to altered biofilm production in sub-inhibitory concentrations of the cell wall-active antibiotics tested above. At 0.5 and 1 µg/mL oxacillin, *prsA*::Tn biofilm production was reduced relative to both OG1RF and Δ*gelE* (Figure 5A). Similarly, *prsA*::Tn biofilm production was reduced relative to Δ*gelE* at 0.5 and 1 µg/mL vancomycin (Figure 5B). This suggests that disruption of *prsA* could affect stability or folding of additional proteins besides GelE that are important for biofilm formation in the presence of these antibiotics. In sub-inhibitory concentrations of penicillin G and ampicillin, Δ*gelE* and *prsA*::Tn had reduced biofilm production relative to OG1RF, but there were no differences in biofilm quantity produced between the 2 mutant strains (Figure 5CD). Because the Δ*bph* mutant has a severe biofilm defect even in the absence of antibiotics (27), it was difficult to interpret any antibiotic-associated changes in biofilm formation for this strain (Figure S4).

**Figure 5.**
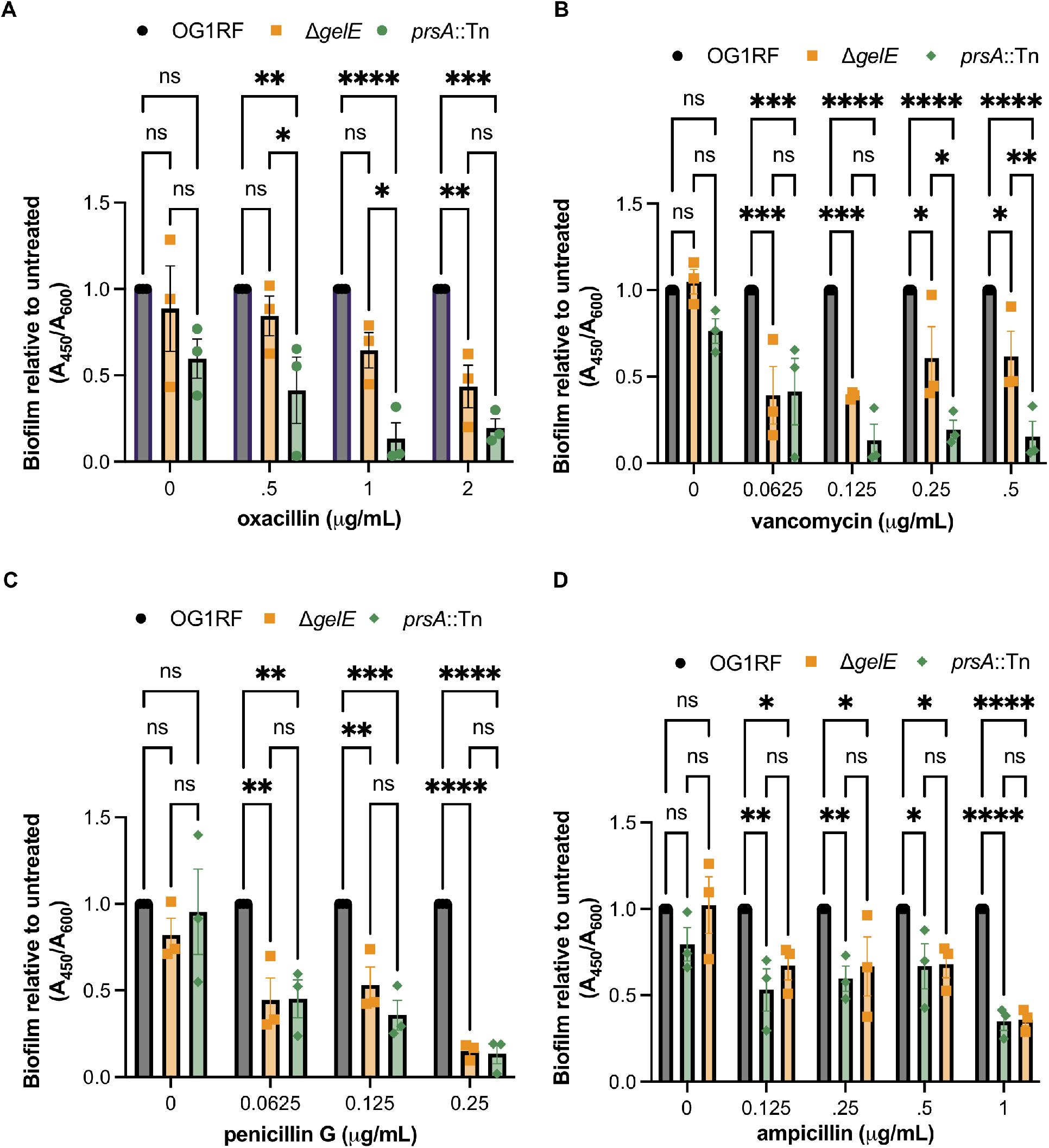
*prsA*::Tn has reduced biofilm production independent of GelE in sub-inhibitory concentrations of oxacillin and vancomycin. The indicated strains were grown in a 2-fold dilution series of **A)** oxacillin, **B)** vancomycin, **C)** penicillin G, and **D)** ampicillin, and growth was measured as A_600_. Biofilm material was stained with safranin and quantified at A_450_. For each mutant, biofilm production was normalized to OG1RF. Data points represent independent biological replicates (n = 3), and error bars show standard error of the mean. *p<0.05, **p<0.01, ***p<0.001, ****p<0.0001 by two-way ANOVA with Tukey’s multiple comparisons test.

### Deletion of *prsA* increases efficiency of conjugative plasmid transfer

GelE affects stability of aggregation substance proteins and pheromone peptides that mediate transfer of conjugative plasmids like pCF10, and plasmid transfer into strains lacking *gelE* is more efficient than into GelE-producing cells (5). We previously showed that the rate of transfer of plasmid pCF10 into Δ*bph* was similar to parental OG1RF (27), suggesting that the reduction in GelE levels in this strain is not enough to affect conjugation. Here, we asked whether the *prsA*::Tn mutation affected plasmid transfer, since this Tn mutant does not produce active gelatinase. Donor cells (OG1Sp pCF10) and recipients (OG1RF, Δ*gelE*, or *prsA*::Tn) were mixed 1:1, and transconjugants were quantified by plating on selective growth medium. As seen previously, transfer to Δ*gelE* recipients was increased ∼2-fold relative to OG1RF. Consistent with the GelE-negative phenotype of *prsA*::Tn, transfer was also increased into these recipients (Figure 6).

**Figure 6.**
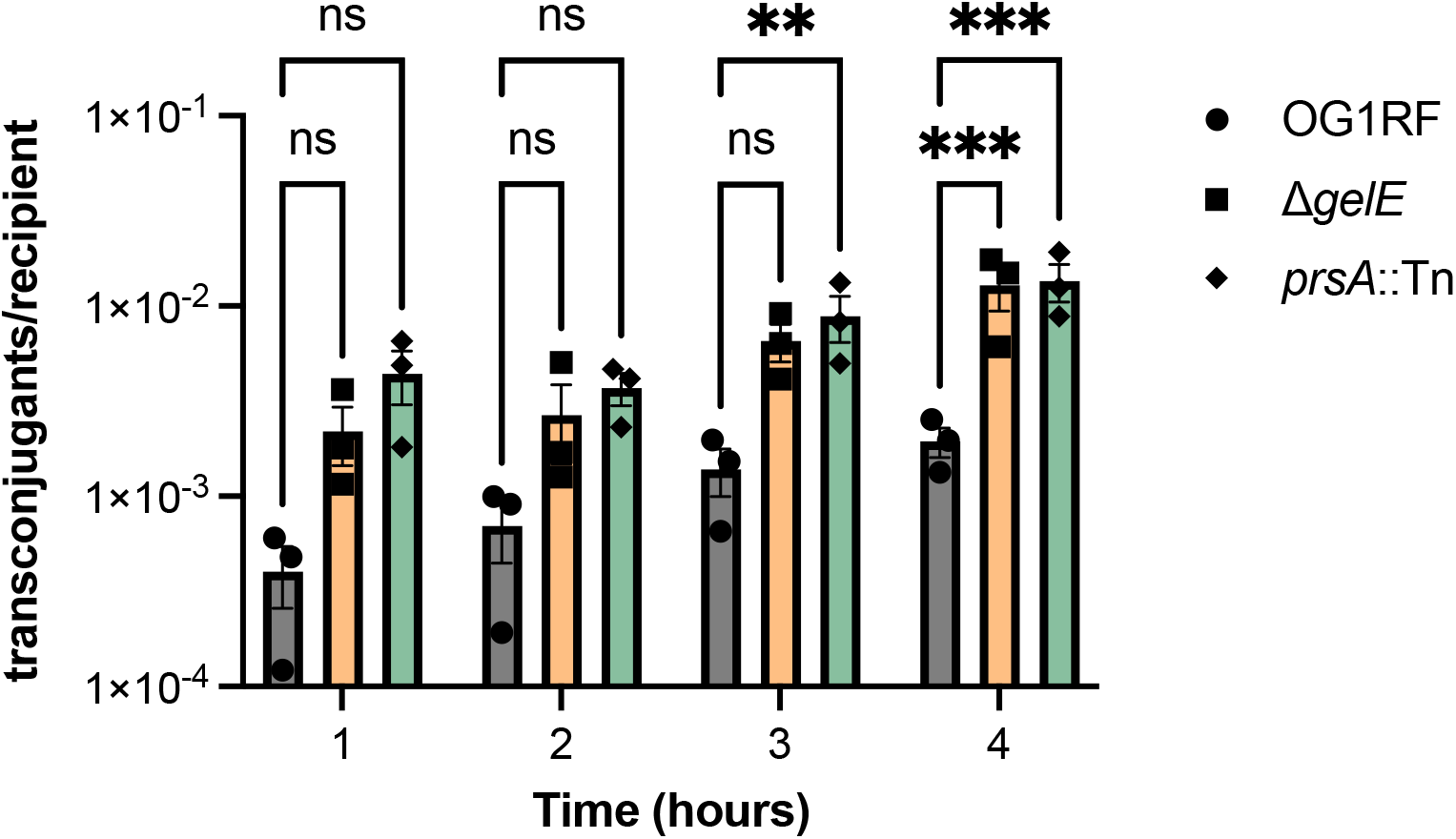
Disruption of *prsA* increases transfer of the conjugative plasmid pCF10. Donors (OG1Sp pCF10) and recipients (OG1RF, Δ*gelE*, or *prsA*::Tn) were mixed 1:1 and incubated at 37 °C. Each data point represents a biological replicate (n = 3). ** p<0.01, *** p<0.0001 by two-way ANOVA with Dunnett’s correction for multiple comparisons.

## Discussion

Gelatinase is an important virulence factor for *E. faecalis*, and regulation of *gelE* expression by the Fsr quorum sensing (QS) system has been extensively studied (12, 13). In 2004, Hancock and Perego proposed there might be another protein required for folding or processing of GelE based on complications purifying active GelE overexpressed in *E. coli* (13). Here, we identify two additional proteins involved in gelatinase production and activity (Figure 7). The biofilm-associated phosphatase Bph regulates *gelE* expression through an Fsr-dependent manner, as expression of the *fsr* locus is reduced in a Δ*bph* mutant. The extracellular peptidyl isomerase PrsA acts upon GelE in an Fsr-independent mechanism, presumably by ensuring proper refolding of GelE as it is secreted from the cell. Interestingly, Δ*bph* has a gelatinase-positive phenotype after overnight growth on gelatin plates (27), although here we showed a significant reduction in gelatinase activity at discrete time points during growth. This highlights the caution that must be taken when interpreting genetic screens that can obscure subtle phenotypes caused by reduced gene expression over a long period of time. Given that our findings update the longstanding model for gelatinase production, it is worth determining whether *E. faecalis* encodes other gene products that regulate gelatinase expression and activity. GelE and Fsr levels increased in an *E. faecalis* mutant lacking the ClpP protease, suggesting that additional factors beyond Fsr, Bph, and PrsA could regulate expression, activity, or stability of gelatinase (44).

**Figure 7.**
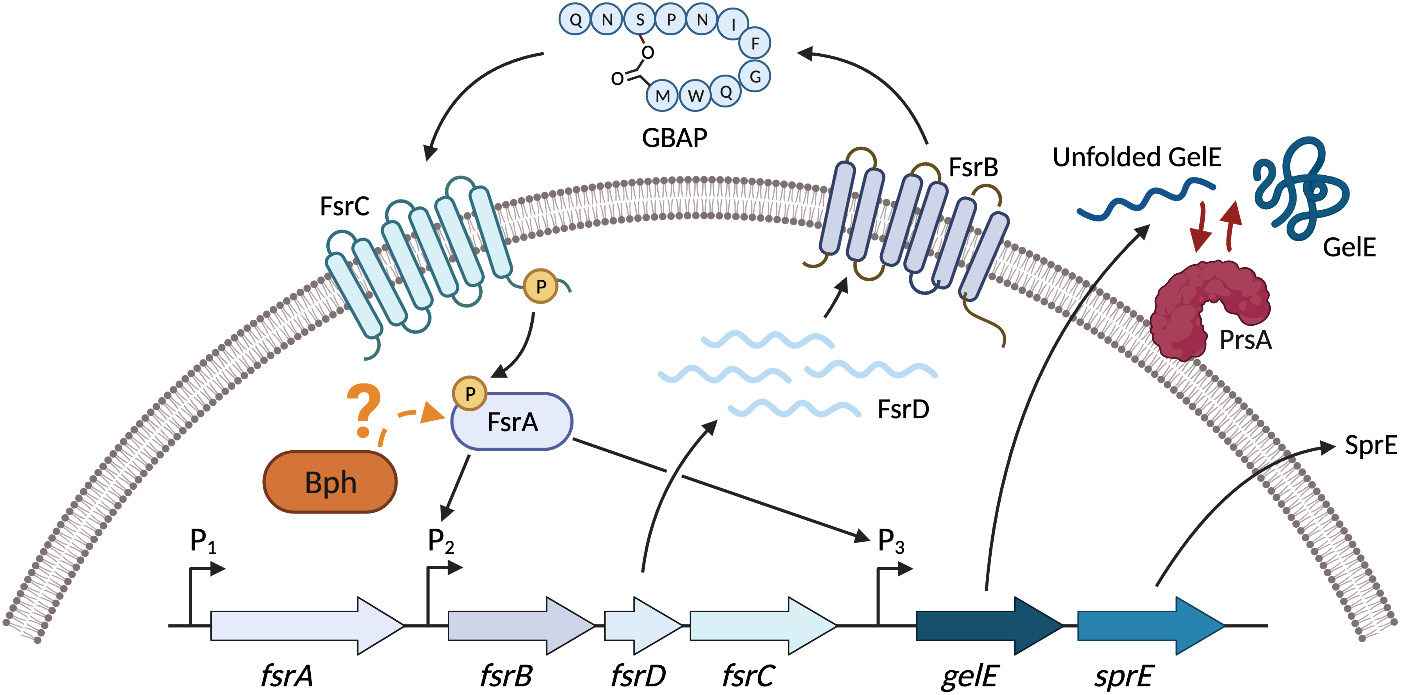
Updated model for gelatinase production in *E. faecalis*. The Fsr quorum sensing system positively regulates expression of the proteases GelE and SprE. Through an unknown mechanism, Bph is required for expression of the *fsr* locus and thus regulation of GelE. The extracellular foldase PrsA is required for activity of secreted GelE, but not *fsr* or *gelE* gene expression. This figure was generated with Biorender.

Although our studies were done *in vitro*, the RNAseq could provide physiological insight into the relevance of Bph *in vivo*. Multiple genes from the ethanolamine utilization (*eut*) operon were upregulated in both *bph* and *fsrB* mutants (31), and the loss of *eut* genes increases *E. faecalis* fitness during gastrointestinal tract (GIT) colonization (45). Therefore, overexpression of this operon may in turn decrease GIT fitness of Δ*bph*. Additionally, genes that contribute to production of the enterococcal polysaccharide surface antigen (Epa) were downregulated, and Epa is required for GIT colonization and persistence (46–49). These results combined with our previous study on Bph activity demonstrate that loss of Bph has pleiotropic effects on quorum sensing, expression of virulence factors, and metabolism (27). However, the target of Bph phosphatase activity is still unknown. Multiple transcription factors are differentially expressed in Δ*bph* (**Supplementary Table 1**), so the overall differences in gene expression may represent the cumulative regulons of these proteins. Bph could potentially directly dephosphorylate transcription factors, but previous work did not identify differential protein phosphorylation using a gel-based approach (27). Dephosphorylation of a small molecule, like nucleotides or second messengers, could cause global changes in gene expression. *fsrBC*, *gelE*, *sprE* are also downregulated in (p)ppGpp-null mutant (50), and recombinant Bph had phosphatase activity on (p)ppGpp *in vitro* (27). Therefore, additional work is required to disentangle the genetic networks disrupted in the Δ*bph* mutant, identify the target of Bph phosphatase activity, and determine the importance of *bph* during colonization and infection.

This study also highlights differences in PrsA function across Gram-positive bacteria. In *B. subtilis*, *prsA* is essential and cannot be deleted *in vitro* without compensatory mutations or increased Mg^2+^ in growth medium (22, 36). Multiple PrsA proteins are produced by *B. anthracis*, *Listeria monocytogenes*, and group A *Streptococcus* (23, 24, 51). Although *E. faecalis* encodes multiple PPIases, it produces only one PrsA homolog, which is not essential for viability. In group A *Streptococcus*, PrsA is required for the proper folding of the secreted exotoxin SpeB (52). The *prsA* gene is encoded adjacent to *speB*, and they are co-transcribed. In *E. faecalis*, *prsA* is in a distal location relative to *gelE*, which is encoded adjacent to the Fsr quorum sensing system that regulates *gelE* expression. Additionally, trigger factor is also required for refolding of SpeB (53, 54), but Tig is not required for GelE activity in OG1RF (35).

Interestingly, neither Δ*bph* nor *prsA*::Tn exactly phenocopied the Δ*gelE* mutant during growth in sub-inhibitory concentrations of vancomycin or β-lactam antibiotics despite the reduced gelatinase levels in those strains. Multiple cell wall biosynthesis genes were downregulated in Δ*bph*, which could affect sensitivity to antibiotics. We also observed protein bands in protoplast, cell wall, and supernatant fractions that were differentially expressed in the *prsA*::Tn mutant relative to OG1RF. This coupled with the biofilm-specific antibiotic sensitivity of *prsA*::Tn suggests that other proteins that sense or respond to antibiotics during biofilm growth could be misfolded in the absence of PrsA. As such, identifying additional targets of PrsA foldase activity (beyond GelE) could serve as a platform for better understanding biofilm-specific antibiotic resistance in *E. faecalis*. We also observed interesting differences in Δ*bph* and *prsA*::Tn antibiotic sensitivity when comparing growth in liquid culture and on agar plates and speculate that this is because growth on an agar plate more closely resembles biofilm growth.

Given the importance of GelE in virulence, it is interesting to consider the potential interplay between *bph*, *prsA*, and gelatinase production during infections. In a rabbit subdermal abscess model, expression of *bph* and *prsA* increased after 2 h infection, whereas *bph* and *gelE* were downregulated after 8 h (55). This suggests that regulation of gelatinase is critical for establishing an infection and *in vivo* survival. Additionally, numerous studies have documented a discrepancy between gelatinase genotype and phenotype in clinical isolates (16, 18, 56–59). However, these collections of isolates are traditionally screened for GelE activity using gelatin plates or tubes, which could obscure temporal changes in gelatinase production as demonstrated in this study. A survey of environmental *Enterococcus* isolates found that GelE-positive *E. faecalis* strains lost the gelatinase phenotype with serial passaging even though *gelE* was still transcribed. (60). Therefore, additional analysis is needed to understand how Bph and PrsA control the dynamics of gelatinase production and how this affects the establishment and persistence of infections in humans.

## Materials and Methods

### Bacterial strains and growth conditions

All strains used in this study are listed in **Supplementary Table 3**. Bacteria were maintained as freezer stocks at −80 °C in 25% glycerol. Strains were cultured in Brain-Heart Infusion broth (BHI), tryptic soy broth without added dextrose (TSB-D), or modified M9 medium supplemented with yeast extract and glucose (MM9-YEG) as indicated. BHI and TSB-D were purchased from BD, and MM9-YEG was prepared as previously described (61). For solid media, agar was added to a final concentration of 1%. Unless indicated otherwise, overnight cultures were grown in the same medium used for experiments. Antibiotics were used at the following concentrations: erythromycin (10 µg/mL), fusidic acid (25 µg/mL), spectinomycin (50 µg/mL for *E. coli*, 250-1000 µg/mL for *E. faecalis*), and tetracycline (5 µg/mL). When needed for induction of gene expression, cCF10 (Mimotopes) or nisin (Sigma) were added to cultures at 25-50 ng/mL or 50 ng/mL, respectively.

### Cloning

Oligonucleotides used for cloning are listed in **Supplementary Table 3**. Plasmids pML200 and pML201 are derivatives of pTCV-LacSpec (62) with the *fsrA* and *fsrB* promoters fused to *lacZ*, respectively. These plasmids were a generous gift from Lynn Hancock and Marta Perego. To obtain pML200, a PCR fragment encompassing ∼1 kb upstream of the *fsrA* gene as well as a portion of the *fsrA* gene was digested with EcoRI and HincII and cloned into pTCV-LacSpec digested with EcoRI and SmaI. To obtain pML201, a PCR fragment encompassing ∼500 bp that included a portion of the *fsrA* gene, the intergenic region, as well as the first 197 bp of the *fsrB* gene was digested with ApoI and HaeIII and cloned into pTCV-LacSpec digested with EcoRI and SmaI. Plasmid pDM7 is a derivative of pTCV-LacSpec plasmid with the *gelE* promoter fused to *lacZ* and was a generous gift from Dawn Manias. To create pheromone-inducible plasmids expressing *fsrA* and *gelE*, alleles were amplified from purified OG1RF genomic DNA using Pfu Ultra II polymerase (Agilent), digested with BamHI-HF and NheI-HF, then ligated to pCIEtm (27) treated with the same restriction enzymes. Restriction enzymes were purchased from New England Biolabs. Plasmid constructs were verified by Sanger sequencing at the University of Minnesota Genomics Center.

### RNAseq

Overnight cultures were diluted to OD600 = 0.05 in TSB-D. At 2 and 4 h, a volume equivalent to OD_600_ = 1 was mixed with an equal volume of RNA protect and incubated at room temperature for 10 min. Cells were pelleted and stored at −80 °C. Pellets from 2 biological replicates were resuspended in 200 µL of buffer (10 mM Tris pH 8.0, 1.0 mM EDTA) supplemented with lysozyme (30 mg/mL) and incubated at 37 °C for 10 min. Total RNA was extracted with a Qiagen RNeasy kit following manufacturer’s instructions and treated with TURBO DNase (Ambion/Thermo Scientific, Waltham, MA) following the manufacturer’s “rigorous treatment” protocol. Removal of contaminating genomic DNA was verified by conventional PCR using 16S primers (JD460s/JD461as, **Supplementary Table 3**). RNA was submitted to the University of Minnesota Genomics Center for ribosomal RNA depletion (Illumina) and TruSeq stranded library preparation. Samples were pooled and sequenced (2×75 bp paired ends) on an Illumina NextSeq 550 in mid-output mode. Sequencing quality was evaluated using FastQC (63). Reads were trimmed with Trimmomatic (64) and imported into Rockhopper for analysis using standard settings (65, 66). Fold changes (log2) were calculated from Rockhopper expression values. Rockhopper-generated q-values ≤ 0.05 were considered statistically significant. For KEGG analysis, identifiers corresponding to OG1RF locus tags were obtained from KEGG and used as input for KEGG Mapper (67, 68).

### qRT-PCR

Overnight cultures were adjusted to OD_600_ = 0.05 in fresh media and grown for 6 h, after which the equivalent of 1 mL cells at OD_600_ = 1.0 was collected. Total RNA isolation, DNase treatment, and confirmation of genomic DNA removal were carried about as described above. An iScript Select cDNA Synthesis Kit (BioRad) was used for cDNA synthesis using random hexamers with 50 µg/mL RNA in a 20 µL reaction. Samples without reverse transcriptase were included to control for DNA contamination. cDNA was diluted 10-fold in sterile water, after which 2 µL were used per qRT-PCR reaction mixture (iQ SYBR Green Supermix, BioRad). Total reaction volumes were 15 µL with 250 nM of each primer (**Supplementary Table 3**). Reactions were run on an iCycler iQ5 (Bio-Rad), and the 2^−ΔΔ*C*T^ method (69) was used to analyze data and calculate the relative fold change normalized to *relA*, which is constitutively expressed in planktonic cultures for each strain.

### Biofilm assays

Cells were grown overnight and diluted 1:100 into fresh TSB-D. 100 µL were added to a 96-well plate (Corning 3595) and incubated in a humidified chamber at 37 °C for 6 or 24 h, as indicated. Overall growth was measured in a Biotek Synergy HT plate reader as absorbance at 600 nm (A_600_). Planktonic cells were removed, and plates were washed gently three times with ultrapure water. Plates were dried overnight in a biosafety cabinet or on the lab bench. Adherent biofilm material was stained with 0.1% safranin (w/v) for 20 minutes at room temperature. Plates were washed and dried as described above. Biofilm material was measured in a Biotek Synergy HT plate reader as A_450_. Biofilm values are presented as safranin-stained material relative to overall cell growth (A_450_/A_600_) normalized to OG1RF. All assays had at least 3 biological replicates, each with technical duplicates.

### Gelatinase and Azocoll assays

Agar plate-based gelatinase assays were performed as described previously (35, 70). Azocoll assays using culture supernatants were modified from published protocols (30, 70). To prepare the substrate, 0.25 g Azocoll (Azo dye-conjugated collagen, Sigma) was washed in 50 mL KPBS (potassium-phosphate buffer saline, pH 7.0) on a rotating platform shaker for 1 h at room temperature, then centrifuged (4,900 × g for 10 min). Azocoll was resuspended, and the wash was repeated. Following the second wash, the residue was resuspended in 50 mL KPBS containing 1 mM CaCl_2_. Washed Azocoll was stored at room temperature and used within 1 week. Strains were grown overnight in TSB-D supplemented with fusidic acid or tetracycline and were diluted to OD_600_ = 0.05 in fresh medium with tetracycline and cCF10 as needed. Cultures were incubated statically at 37 °C. At each time point, OD_600_ was measured, and 1.5 mL cells were pelleted in a tabletop centrifuge (6,080 × g for 2 min). 500 µL supernatant was mixed with 500 µL washed Azocoll in a 1.5 mL tube, and tubes were incubated at 37 °C on a shaker for 24 h. Samples were centrifuged (16,000 × g for 5 min), and dye release was quantified in a Biotek Synergy HT plate reader as absorbance at 550 nm (A_550_). Data is expressed as dye release relative to growth (A_550_/OD_600_) from three biological replicates with technical replicates (3 per sample). Due to variations between batches of Azocoll, all replicates for a given experiment were done using Azocoll prepared from the same bottle.

### β-galactosidase activity assays

Strains were grown overnight in BHI containing spectinomycin. For blue-white plate screening, 5 µL of each overnight culture was spotted on MM9-YEG plates containing spectinomycin and 200 µg/mL X-Gal (5-bromo-4-chloro-3-indolyl-β-d-galactopyranoside). Plates were incubated overnight at 37 °C and photographed with a cell phone camera. To quantify β-galactosidase activity, overnight cultures were diluted to OD_600_ = 0.05 in MM9-YEG and spectinomycin and incubated for 4 h at 37 °C. The cells were pelleted (6,080 × g for 2 min) and stored at −80 °C. β-galactosidase expression was measured as previously described (71). Assays were performed in biological triplicate with technical duplicates.

### SDS-PAGE

Overnight cultures grown in either TSB-D or MM9-YEG (with tetracycline and cCF10, as needed) were diluted to OD_600_ = 0.05 in fresh media. After 6 hr static incubation at 37 °C, the equivalent of 1 mL cells at OD_600_ = 1.0 was pelleted in a tabletop centrifuge (6,080 × g) for 2 min. Pellets were resuspended in 100 µL TESL (10 mM Tris-HCl pH 8.0, 1 mM EDTA, 25% sucrose, and 15 mg/mL lysozyme) and incubated at 37 °C for 30 min. 15 µL was mixed directly with 2× Laemmli sample buffer (BioRad) for whole-cell lysates, and 85 µL was centrifuged (16,000 × g) for 1 min to separate the pellet (protoplast) from the supernatant (cell wall extract). Cell wall extracts were mixed with an equal volume of 2× Laemmli sample buffer. Protoplast samples were resuspended in 50 uL urea lysis buffer (8 M urea, 20 mM Tris–HCl pH 7.5, 150 mM NaCl) and 50 uL 2× Laemmli sample buffer. Proteins in the culture supernatant were precipitated by mixing 1 volume of supernatant with 0.25 volumes of chilled 100% trichloroacetic acid on ice, pelleted in a tabletop centrifuge (16,000 × g) for 10 min, and washed twice with acetone. Pellets were dried and resuspended in 50 uL urea lysis buffer and 50 uL 2× Laemmli sample buffer. Samples were heated at 95 °C before loading onto 10% Tris-glycine SDS-PAGE gels. Gels were run at 110V, stained with Coomassie, destained, and imaged on a BioRad Gel Doc EZ Imager.

### Antibiotic sensitivity assays

Overnight cultures were grown in TSB-D and adjusted to 5 ×10^7^ CFU/mL in fresh medium. Cells were diluted 1:50 into 96-well plates (Corning 3595) containing 100 µL of two-fold serial dilutions of the indicated antibiotics prepared in TSB-D. Plates were incubated in a humidified chamber at 37 °C for 24 h after which OD_600_ measurements were taken in a BioTek Synergy H1 plate reader. Relative growth was quantified by dividing the OD_600_ at each antibiotic concentration by the OD_600_ of the untreated culture. The results shown are the mean OD_600_ values from three independent biological replicates with technical duplicates. To measure biofilm production in the presence of sub-inhibitory antibiotic concentrations, the 96-well plates used for antibiotic sensitivity assays were dried and stained with safranin as described above.

### Statistical analysis

Data analysis was performed using GraphPad Prism (version 9.2.0). Statistical tests and significance thresholds are described in the figure legends.

## Data availability

Files generated from RNA sequencing and data analysis have been deposited with NCBI GEO under accession number GSE198051.

## Acknowledgements

We thank Lynn Hancock, Marta Perego, and Dawn Manias for providing plasmid constructs. The authors acknowledge the Minnesota Supercomputing Institute (MSI) at the University of Minnesota (http://www.msi.umn.edu) and the University of Minnesota Genomics Center (https://genomics.umn.edu/) for providing resources that contributed to the results reported here. This work was supported by the National Institutes of Health grants 1R35GM11807 and 1RO1AI122742 to GMD, 1K99AI151080 to JLEW, and the National Center for Advancing Translational Sciences grant UL1TR002494. EBR was supported by the University of Minnesota Undergraduate Research Opportunities Program (UROP). The content is solely the responsibility of the authors and does not necessarily represent the official views of the National Institutes of Health’s National Center for Advancing Translational Sciences.

---

**Figure S1.**
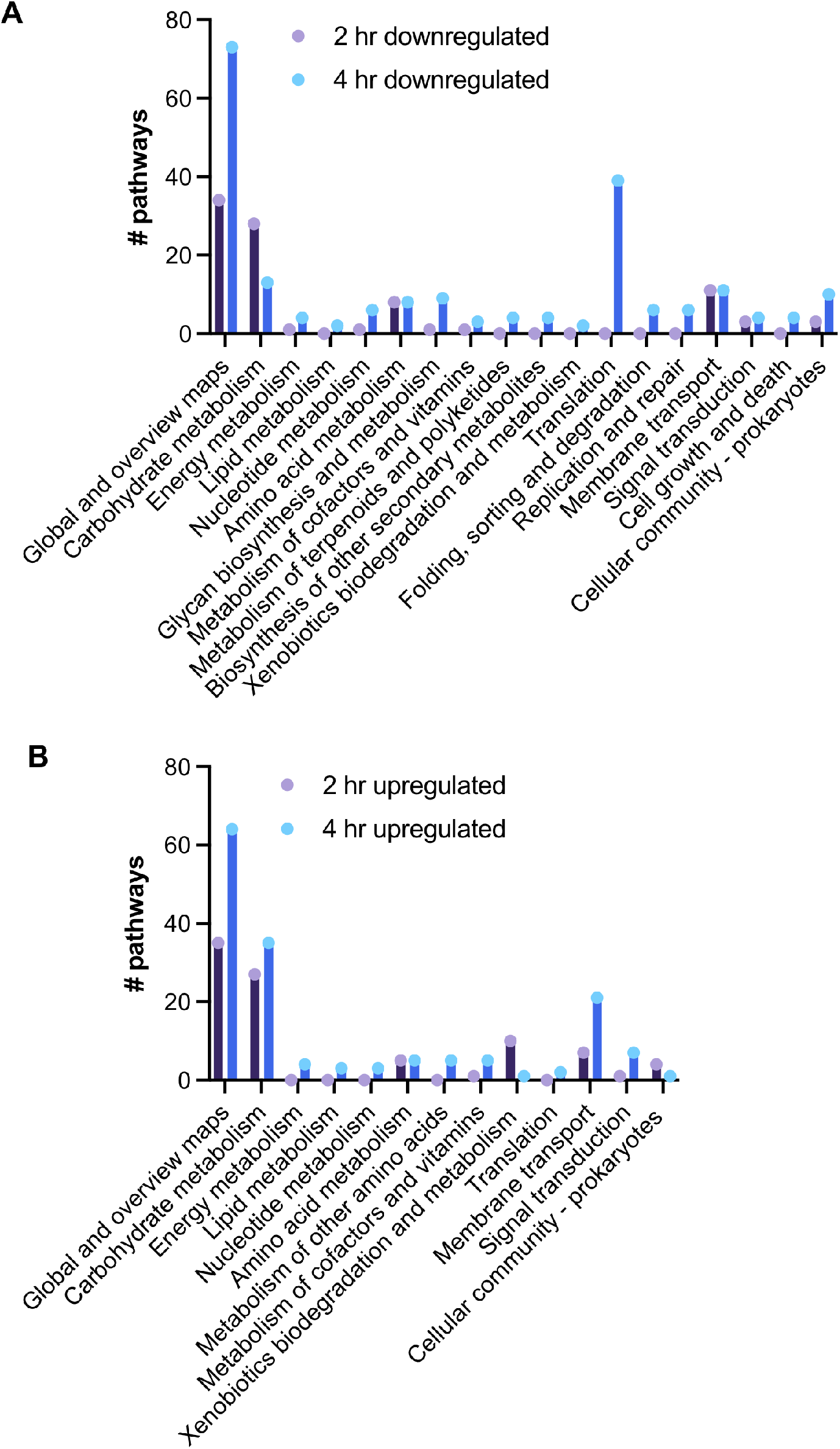
Categories of differentially expressed genes in Δ*bph* after 2 and 4 h planktonic growth. OG1RF locus tags were converted to KEGG identifiers, and category analysis was done using KEGG Mapper for **A)** downregulated and **B)** upregulated genes.

**Figure S2:**
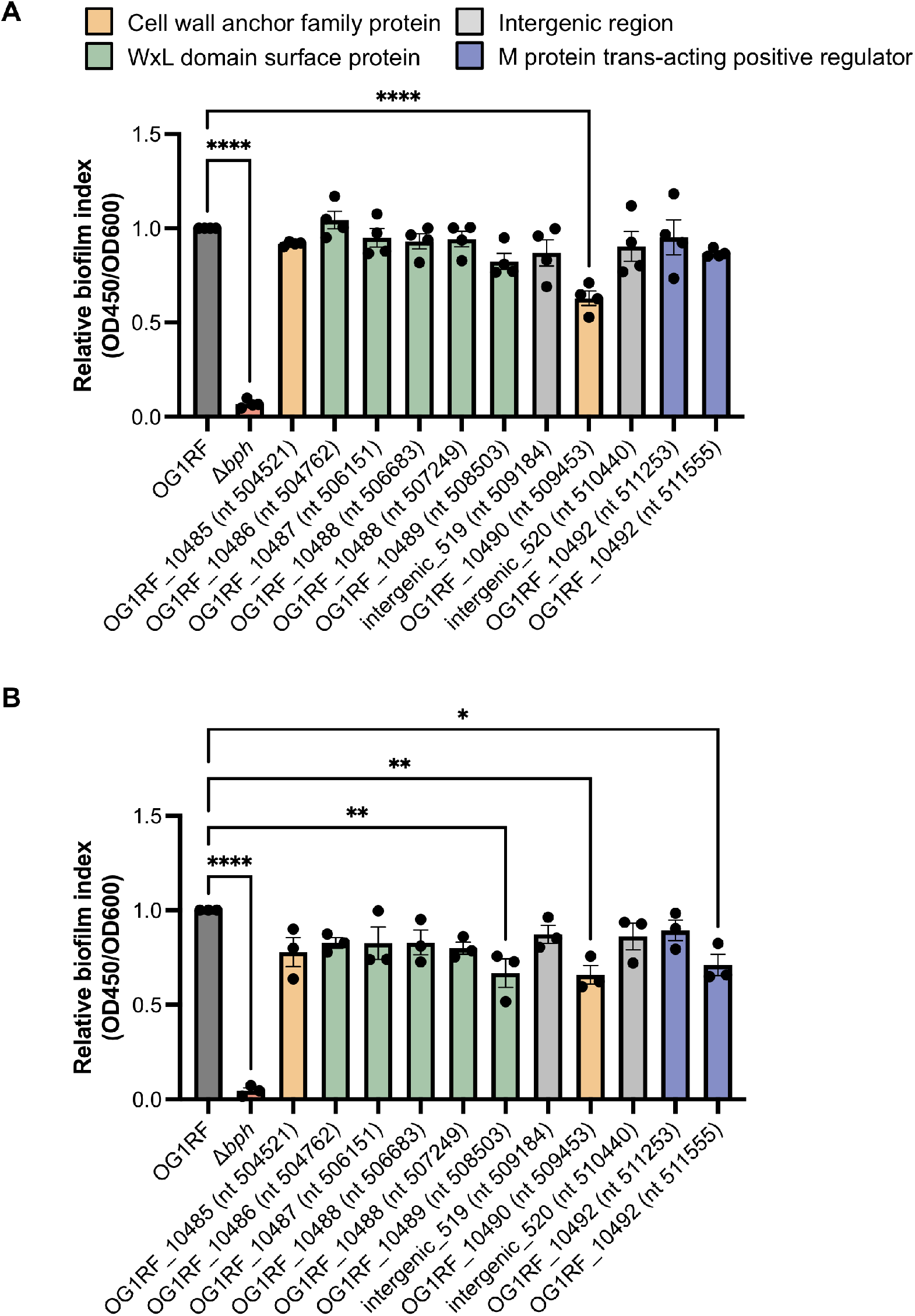
Biofilm formation of mutants with Tn insertions in OG1RF_10485-10492. Tn mutants were obtained from stock Tn library plates, and biofilm formation in TSB-D was tested in 96-well plates at **A)** 6 h and **B)** 24 h. Biofilm biomass was detected by safranin staining (A_450_), and biofilm production was calculated relative to overall growth (A_600_). Values were normalized to OG1RF. Each data point represents an independent biological replicate (**A**, n = 4; **B**, n = 3). Error bars represent standard error of the mean. Statistical significance was calculated by one-way ANOVA (*p<0.05, **p<0.01, ****p<0.0001).

**Figure S3.**
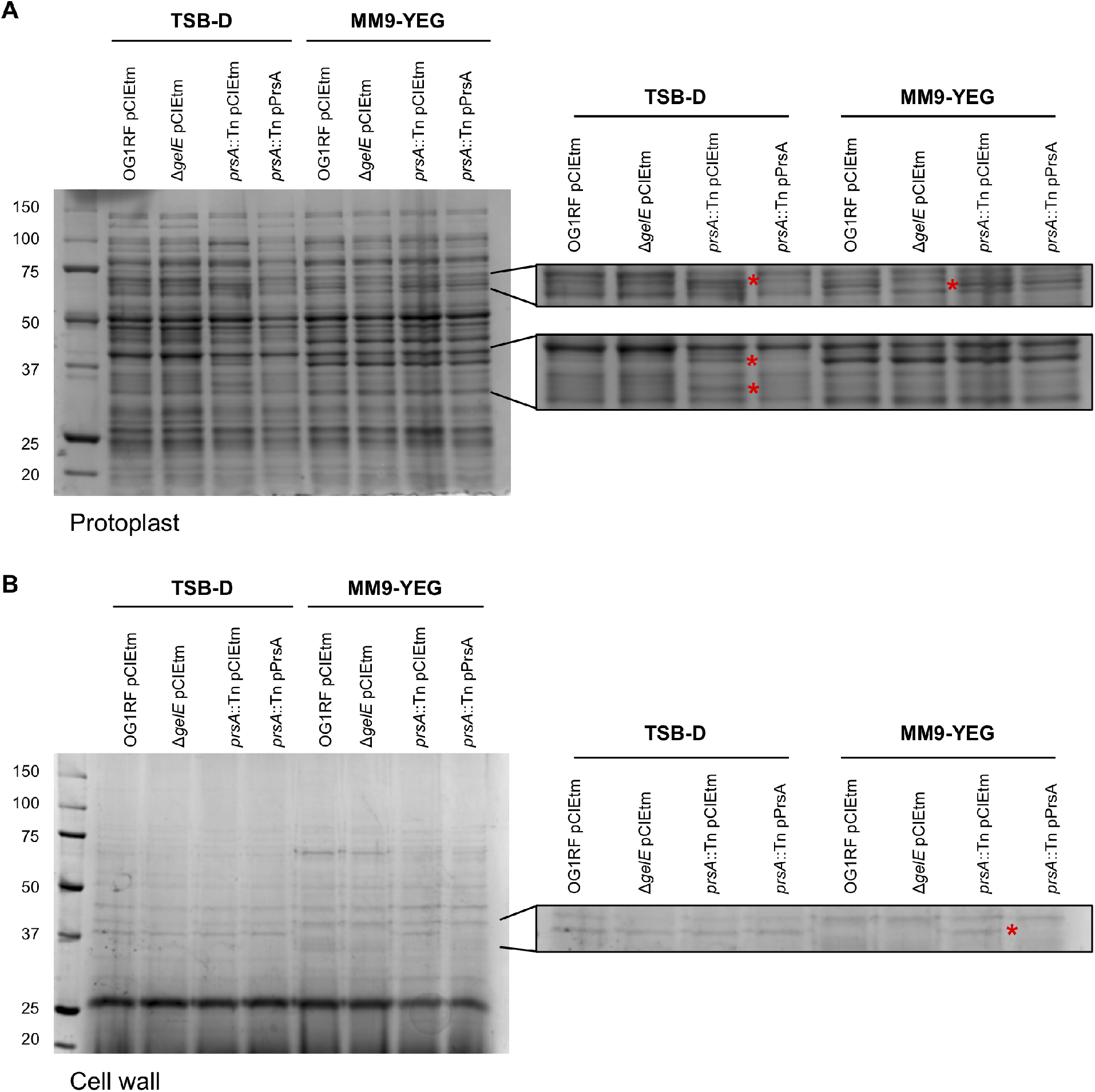
Protein expression in OG1RF, Δ*gelE*, and *prsA*::Tn. Protein lysates were prepared from **A)** protoplasts and **B**) cell wall samples. Samples were run on SDS-PAGE gels and visualized via Coomassie staining. Protein bands differentially expressed in mutant strains are marked with a red asterisk to the right of the band. Images are representative of three biological replicates.

**Figure S4.**
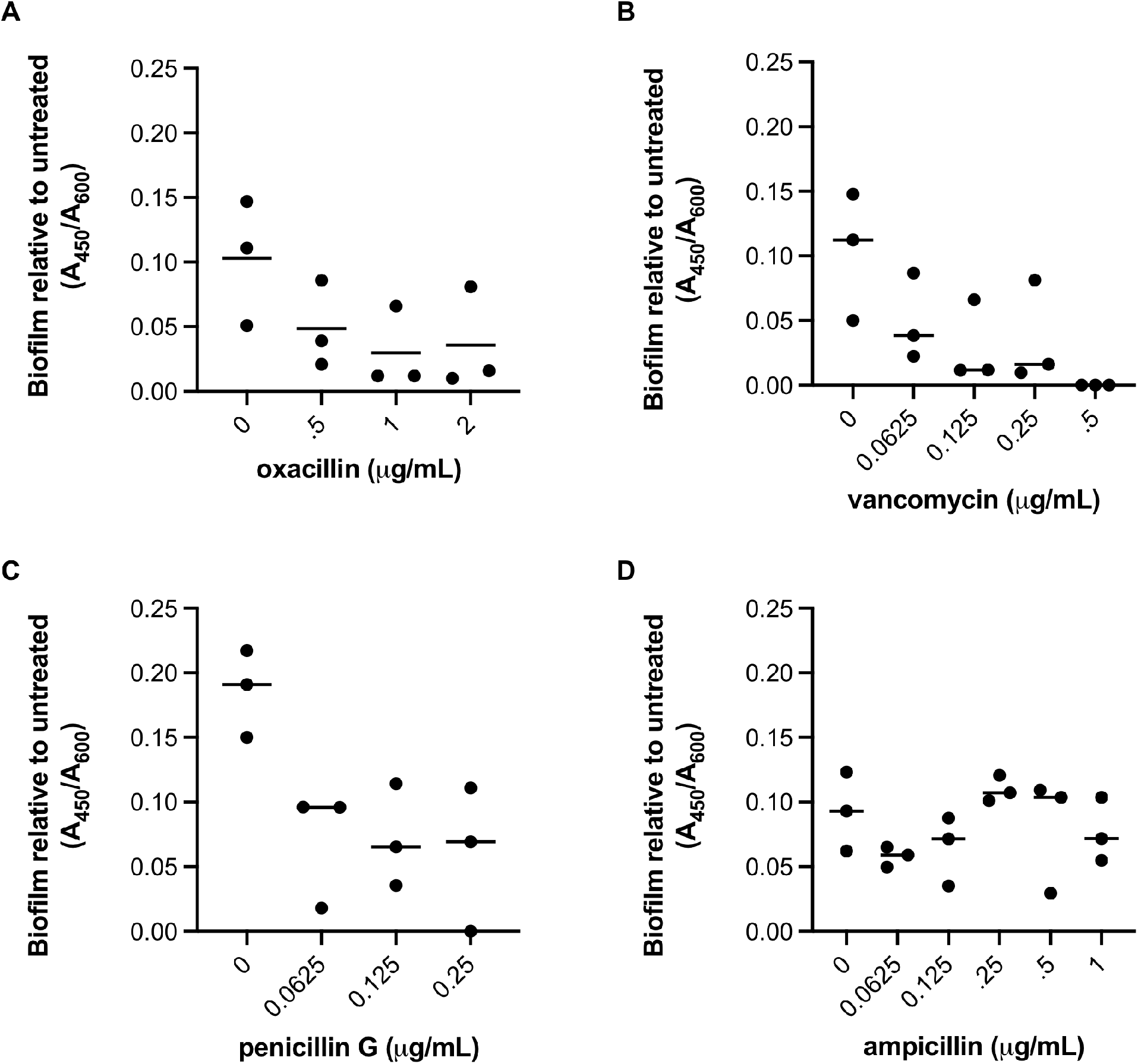
Biofilm production of Δ*bph* in the presence of sub-inhibitory concentrations of antibiotics. Δ*bph* was grown in a 2-fold dilution series of **A)** oxacillin, **B)** vancomycin, **C)** penicillin G, and **D)** ampicillin, and growth was measured as A_600_. Biofilm material was stained with safranin and quantified at A_450_. Biofilm production was calculated relative to OG1RF for each sample. Data points represent independent biological replicates (n = 3), and error bars show standard error of the mean.

**Supplementary Table 1: RNAseq analysis of genes differentially expressed in Δ*bph* compared to OG1RF at 2 and 4 hrs in planktonic culture.**

**Supplementary Table 2. Comparison of differentially expressed genes in Δ*bph* and Δ*fsrB* deletion mutants.**

